# The role of LHCBM1 in non-photochemical quenching in *Chlamydomonas reinhardtii*

**DOI:** 10.1101/2022.01.13.476201

**Authors:** Xin Liu, Wojciech Nawrocki, Roberta Croce

## Abstract

Non-photochemical quenching (NPQ) is the process that protects photosynthetic organisms from photodamage by dissipating the energy absorbed in excess as heat. In the model green alga *Chlamydomonas reinhardtii*, NPQ was abolished in the knock-out mutants of the pigment-protein complexes LHCSR3 and LHCBM1. However, while LHCSR3 was shown to be a pH sensor and switching to a quenched conformation at low pH, the role of LHCBM1 in NPQ has not been elucidated yet. In this work, we combine biochemical and physiological measurements to study short-term high light acclimation of *npq5*, the mutant lacking LHCBM1. We show that while in low light in the absence of this complex, the antenna size of PSII is smaller than in its presence, this effect is marginal in high light, implying that a reduction of the antenna is not responsible for the low NPQ. We also show that the mutant expresses LHCSR3 at the WT level in high light, indicating that the absence of this complex is also not the reason. Finally, NPQ remains low in the mutant even when the pH is artificially lowered to values that can switch LHCSR3 to the quenched conformation. It is concluded that both LHCSR3 and LHCBM1 need to be present for the induction of NPQ and that LHCBM1 is the interacting partner of LHCSR3. This interaction can either enhance the quenching capacity of LHCSR3 or connect this complex with the PSII supercomplex.

## Introduction

All photosynthetic organisms need to deal with constantly changing environmental factors. Changes in light intensity are particularly critical since light is the source of energy but can also become the source of damage. In low light, most of the photons absorbed by the pigments associated with the photosynthetic complexes lead to charge separation in the reaction centers of Photosystem I and II. In high light, when the capacity of the photosynthetic reactions is saturated, the photons absorbed in excess can lead to PSII photoinhibition (Tyystjarvi, 2013; Nawrocki et al., 2021) and the production of harmful species that can damage the photosynthetic apparatus (Erickson et al., 2015; Li et al., 2018). To adapt to the rapid light changes, all photosynthetic organisms have developed a range of photoprotection mechanisms.

A quickly activated photoprotection mechanism, called non-photochemical quenching (NPQ) dissipates as heat a large part of the energy absorbed in excess (Ruban, 2016, 2018). In the model green alga *Chlamydomon*as *reinhardtii*, NPQ activation depends on the presence of the Light-harvesting stress-related complex 3 (LHCSR3) (Peers et al., 2009). This complex is expressed only upon exposure of the cells to high light in photoautotrophic growth conditions (Allorent et al., 2013; Polukhina et al., 2016). At variance with PsbS, the central protein for NPQ in plants (Li et al., 2000; Fan et al., 2015), LHCSR3 binds pigments and switches from a light-harvesting to a quenched state upon protonation of its lumen-exposed residues (Bonente et al., 2011; Liguori et al., 2013). The switch is triggered by the acidification of the lumen driven by photosynthetic electron transfer. LHCSR3 has thus been proposed to act both as pH sensor and quencher (Bonente et al., 2011; Tian et al., 2019). In addition to LHCSR3, another stress-related protein, LHCSR1, was also shown to induce quenching at low pH (Dinc et al., 2016).

The LHCSRs are members of the light-harvesting complex (LHC) multigenic family, containing all the peripheral antenna complexes of green algae and plants (Engelken et al., 2010; Niyogi and Truong, 2013). These integral membrane proteins accommodate up to 18 pigments (chlorophylls *a* and *b* and carotenoids) and serve as an antenna of both Photosystem I (PSI) and II (PSII) (Croce and van Amerongen, 2020; Pan et al., 2020). In *C. reinhardtii*, the major light-harvesting complexes (LHCBMs) are encoded by the *lhcbm1*-*lhcbm9* genes (Merchant et al., 2007). The nine LHCBM complexes are divided into four subgroups according to their sequence homology: type I (LHCBM3/4/6/8/9), type II (LHCBM5), type III (LHCBM2/7) and type IV (LHCBM1) (Minagawa and Takahashi, 2004).

The individual LHCBMs have similar biochemical and biophysical properties (Natali and Croce, 2015) but were suggested to have different roles (Takahashi et al., 2006; Nguyen et al., 2008; Drop et al., 2014; Drop et al., 2014; Grewe et al., 2014; Girolomoni et al., 2017). In particular, LHCBM1 was shown to be involved in photoprotection since the *C. reinhardtii npq5* mutant, deficient in this subunit, exhibits low NPQ (Elrad et al., 2002; Ferrante et al., 2012). However, the role of LHCBM1 in NPQ is at present unknown. In this work, we use biochemical and physiological approaches to investigate the factors hampering the NPQ development in the absence of LHCBM1, analyzing the photoprotection capacities and photosynthetic properties in *npq5* during high light adaptation.

## Methods

### Strains and growth conditions

The null LHCBM1 mutant *npq5* (Elrad et al., 2002) (alternative name CC-4073) was used in this work. WT (CC-425), the parental strain used for insertional mutagenesis to generate *npq5* (Niyogi et al., 1997), was used as the reference strain. *npq4* (Peers et al., 2009) which lacks both *lhcsr3*.*1* and *lhcsr3*.*2* genes is not able to perform the fast energy-dependent quenching and was used as a negative control. *lhcsr1* mutant (Allorent et al., 2016) and its control 4A+ were used to test the impact of LHCSR1 knock-out on NPQ. All the strains were grown in liquid TAP medium (Gorman and Levine, 1965) (25 °C 140 rpm/min) at least 72h in low light (LL, less than 15 μmol photons*m^-2^*s^-1^) to reach the exponential phase. One day before the HL exposure, the strains were diluted in high salt medium (HSM) (Sueoka, 1960) in LL. On the day of the experiment, the cells were harvested by centrifugation and resuspended in fresh HSM. All cells were adjusted to OD_750_ 0.2 and incubated in LL for 1h before high light (HL, 500 μmol photons*m^-2^*s^-1^) treatment. Aliquots were collected at different time points: 0h HL (1h LL), 24h HL and 48h HL. In the case of CC-425, 50 mg/L arginine was added to the media.

### In vivo photosynthetic measurements

#### NPQ

To quantify light-induced NPQ, a short 5.5-minute protocol was used, in which the illumination phase was 2.5 minutes after a 10-second dark period (when F_0_ was acquired), and the subsequent recovery period (in darkness) was 3 minutes. NPQ was calculated as (F_m_/F_m_’)-1, and the first 50-second fluorescence signals (during illumination) were used for maximum NPQ calculation to avoid the possible superposition of state transitions that occur on longer timescales. The maximum quantum efficiency of PSII photochemistry (F_v_/F_m_) was calculated as (F_m_-F_0_)/F_m_ (Baker, 2008). Actinic light 1500 μmol photons*m^-2^*s^-1^, saturating pulses of >10000 μmol photons*m^-2^*s^-1^ and 180 ms pulse duration were used. All actinic light was red, peaking at 630 nm. Weak detecting light flashes were obtained by filtering broad white light LEDs with 10 mm interference Schott filter (520 nm, 10 nm FWHM).

Acid-induced NPQ was performed as previously reported (Tian et al., 2019). In brief, 1 M acetic acid was added to the culture to decrease the pH to 5.5, and 2 M KOH was used to set the pH to 7 (Tian et al., 2019). Acid-induced quenching ((F_m_/F_m_’)-1) was recorded in darkness using a DUAL-PAM 100 (Walz, Germany) with saturating pulses of 12000 μmol photons*m^-2^*s^-1^ and 180 ms duration. Cells were dark-adapted for at least 30 min before measurements. To measure light-induced quenching, a short actinic light period was also used (∼1500 μmol photons*m^-2^*s^-1^).

#### Maximal PSII rate

The fluorescence rise upon transition of dark-adapted cells (Q_A_ maximally oxidized) to light (Q_A_ maximally reduced) in the presence 10 uM dichlorobenzyl dimethyl urea (DCMU) was measured using 630 nm, 35 μmol photons*m^-2^*s^-1^ actinic light according to (Lazar, 1999) with minor modifications (Nawrocki et al., 2019; Tian et al., 2019).

#### PSII turnover rate in continuous light

The PSII turnover rate was measured with a JTS-10 spectrophotometer according to (Lazar, 1999; Nawrocki et al., 2019) with minor modifications. PSII turnover rate was calculated as: [(PSII operating efficiency, Y(II)) * (the extrapolated maximal PSII rate))] at the same light intensity. Y(II) was measured via a light curve in a DUAL-PAM 100 at 13, 37, 70, 166, 273, 430, 660 and 1028 μmol photons*m^-2^*s^-1^. The maximal PSII rate at 35 μmol photons*m^-2^*s^-1^ was measured in a JTS-10 (see above). A linear fit forced through zero and the value at 35 μmol photons*m^-2^*s^-1^ was then used to extrapolate the maximal rate to each intensity used in the PAM for Y(II) measurements. Cells were kept in darkness for at least 30 min before the experiments.

#### Electrochromic shift (ECS)

The functional PSI/PSII (RC:RC) ratio was measured using single-turnover laser flashes in vivo according to (Joliot and Joliot, 2005; Alric et al., 2010) with the modifications described in (Nawrocki et al., 2016).

### Protein analysis

Total cell protein extracts and immunoblots were performed according to (Ramundo et al., 2013; Dinc et al., 2014). 5ug total cell protein extracts were loaded on each well of stacking gel. All antibodies were purchased from Agrisera (Sweden): LHCBM5 (AS09 408), LHCSR1 (AS14 2819), LHCSR3 (AS14 2766), CP47 (AS04 038), PsaA (AS06 172), ATPC (AS08 312), Cyt f (AS06 119), and Rubisco Larger (AS03 037).

## Results

To confirm that *npq5* is an antenna mutant, we compared the antenna composition in low light-grown cells (LL) of WT (CC-425) and *npq5* (Fig. 1). LHCBM1 was undetectable in *npq5* in LL (Fig. 1A), in agreement with previous results (Elrad et al., 2002). The LHCBMs/CP47 protein ratio in *npq5* was ∼20% lower than in WT (CC-425) (Fig. 1B). This difference is mainly due to the absence of LHCBM1, since the level of the other LHCBMs is similar to that of WT (CC-425) (Figure S1 and S2). The absence of LHCBM1 in the mutant is thus not compensated by an increase of other LHCBMs.

**Fig. 1.**
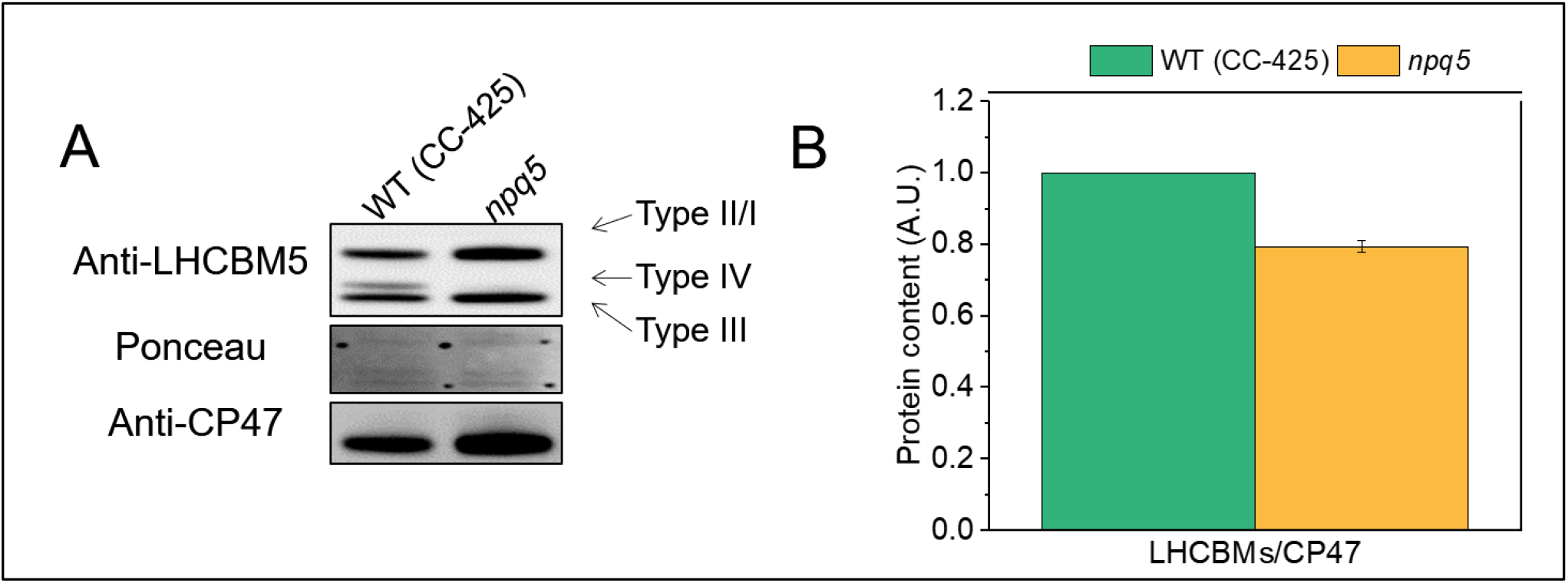
PSII antenna quantification in LL grown cells. Immunoblot with anti-LHCBM5 and -CP47 antibodies (A) and protein quantification (B) are shown for cells grown in low light (<15 μmol photons*m^-2^*s^-1^, 0h HL). The different types (I to IV) of LHCBMs are indicated in (A). Note that the anti-LHCBM5 antibody reacts with all types of LHCBMs. Densitometry data of each protein were normalized to the WT (CC-425) before calculating the ratio in (B). Data shown are mean ± SEM, n=3 biological replicas.

To study the role of LHCBM1 in photoprotection, we measured the photosynthetic properties of the *npq5* mutant and its control strain (WT CC-425) during HL exposure. The cells pre-grown in TAP in low light (LL, 20 μmol photons*m^-2^*s^-1^) were resuspended in HSM, kept for 1h in LL (0h HL) and then exposed to high light (HL, 500 μmol photons*m^-2^*s^-1^). Aliquots were collected at 0h HL, 24h HL and 48h HL for the analysis.

### PSII antenna size and composition upon high light acclimation

Several parameters were measured before and during the HL treatment. The maximum quantum efficiency of PSII (F_v_/F_m_) decreased significantly during HL, suggesting that the cells were partially photoinhibited (Fig. 2A). The F_v_/F_m_ value of *npq5* was identical to its control in LL and after 24h of HL and only slightly lower after 48h (Fig. 2A). These data indicate that *npq5* is relatively well protected in high light, although less than its parental strain in the long run.

**Fig. 2.**
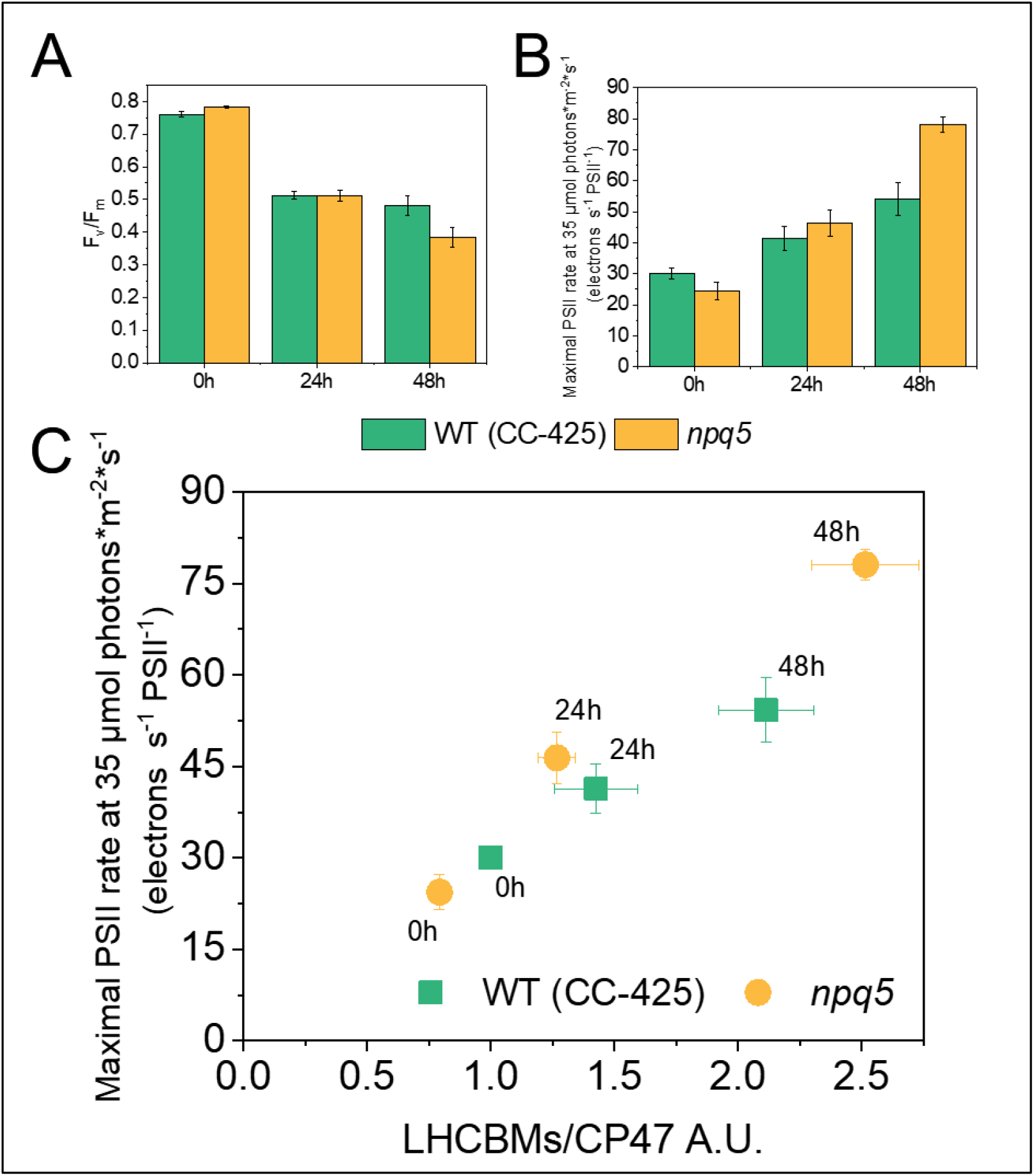
PSII’s efficiency and the antenna size. (A) Maximum PSII quantum efficiency (F_v_/F_m_); (B) functional PSII antenna size; (C) LHCBMs/CP47 protein ratio vs functional PSII antenna size. Densitometry data of LHCBMs and CP47 were each normalized in WT (CC-425) at 0h before calculating the ratio in (C). Data shown are mean ± SEM, n=3 biological replicas. The actinic light source peaks at 630 nm.

Since a change of antenna size is a well-known response to HL exposure (Neale and Melis, 1986; Croce, 2020), the functional antenna size of PSII was measured. Before HL treatment (i.e. in LL), the functional antenna size of PSII was around 20% smaller in *npq5* than in WT (CC-425) (Fig. 2B). This value matches the protein data (Fig. 1B), indicating that the absence of LHCBM1 does not influence the functional organization of the other antennae. Upon HL exposure, the functional PSII antenna size increased in both strains and after 48h HL was significantly higher in *npq5* than in WT (CC-425) (Fig. 2B). Immunoblots against all LHCBM types showed that the relative amount of each subtype remained unchanged in WT (CC-425) and *npq5* during HL exposure (Fig. S1), indicating that there is no specific increase of any of the subunits. Thus, the increase in antenna size upon HL exposure is likely due to a loss of functional core complexes, which results in a larger antenna for the remaining functional ones due to the energetic connectivity (P. Joliot, 1964).

The PSII functional antenna size(Fig. 2C, S2) correlates well with the LHCBMs/CP47 protein ratio in both strains after 24h of HL, while there is some deviation upon 48h. These results suggest that while upon 24h HL most antenna proteins are functionally connected to the core, after longer HL exposure part of the antenna is not well connected anymore.

### Electron transport is impaired in high light in *npq5*

Next, we checked possible differences between WT and mutant in the components and functionality of the electron transport chain. To calculate the PSI/PSII ratio, we quantified the proteins using immunoblots and the charge separation by measuring the electrochromic shift (ECS). *npq5* cells grown in LL showed a slightly lower PSI/PSII ratio than the control strain. This agrees with previous results, which demonstrated that a decrease in PSII antenna size is compensated by a change in PSI/PSII ratio (Nicol et al., 2019; Xu et al., 2020). The agreement between the functional and the protein measurements indicates that all photosystems present in the membrane work properly. The functional data showed a significant increase in the PSI/PSII ratio during HL treatment, especially in *npq5* after 48h HL (Fig. 3A). On the contrary, at the protein level, a decrease in PSI/PSII ratio was observed (Fig. 3B). These results suggest that part of PSII is not functional in high light and that this population is larger in *npq5* than in the WT (CC-425). The cells respond to the reduced PSII functionality by lowering the amount of PSI to restore a balance between the two photosystems.

**Fig. 3.**
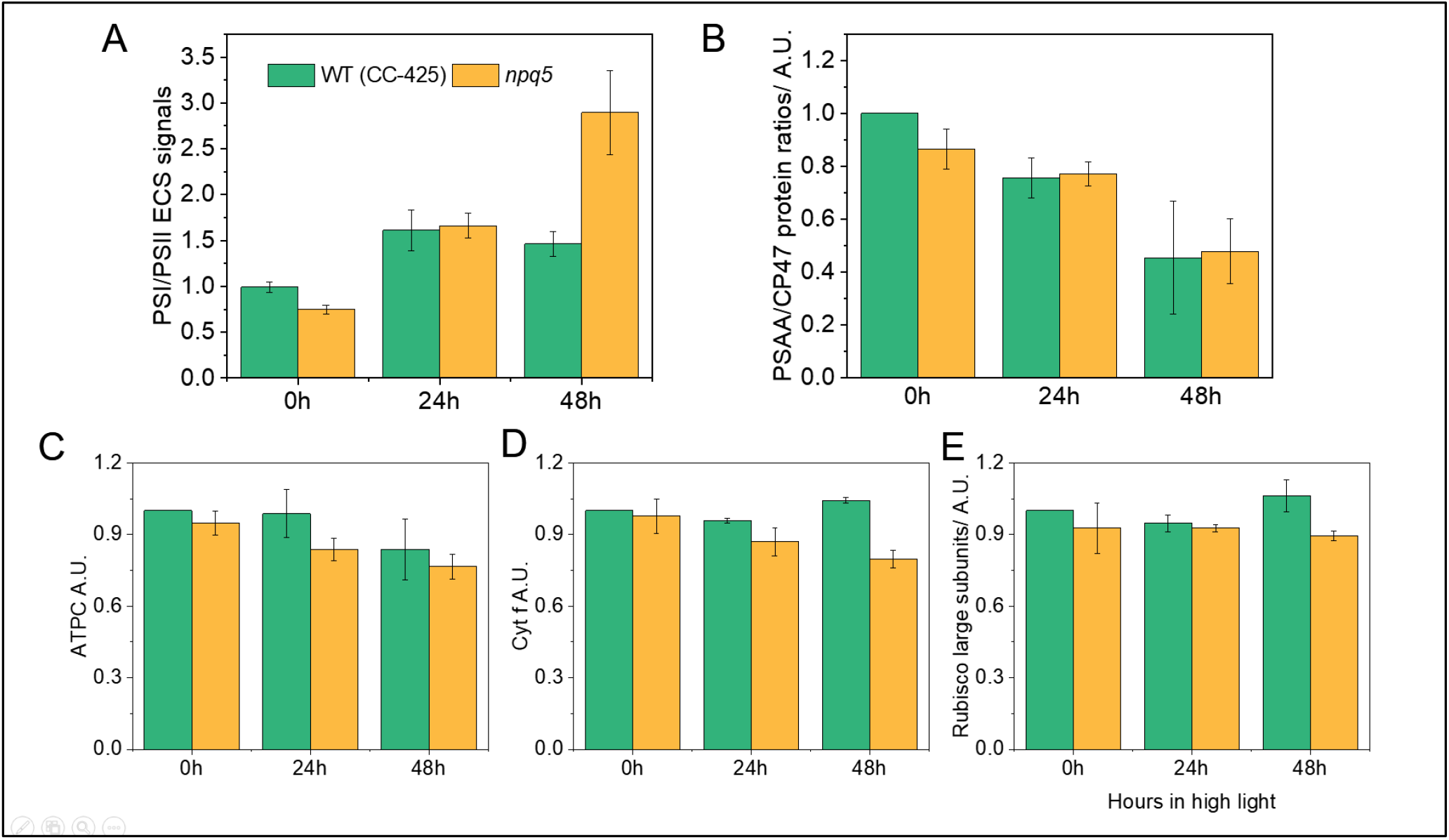
Protein composition of photosynthetic apparatus and PSI/PSII ratios during HL acclimation. The functional measurement of PSI/PSII ratio are based on the ECS signal (A), PSAA/CP47 protein ratio (B), the amount of ATP gamma subunit (ATPC) (C), cytochrome f (Cyt f) (D) and Rubisco Large subunits (E) are shown. Data shown were normalized to WT (CC-425) at 0h in (B-E) and are mean ± SEM, n=3 biological replicas with each of 1 technical replica (B-E) and 4 technical replicas (A). 5 μg total protein extracts were loaded.

To check if the electron transport rate of PSII is affected in *npq5*, we measured PSII turnover rate in continuous light during HL treatment by multiplying the Y(II) parameter by the cross-section of PSII at a given light intensity (Lazar, 1999; Nawrocki et al., 2019) (Fig. 4, S3). The results show that the turnover rate (per PSII) in *npq5* is lower than in WT (CC-425) in all light conditions and that it further decreases upon 48h HL. This is consistent with partial PSII photoinhibition in the mutant, and shows that the ETR is limited by factors other than the absorption cross-section, which increased under HL treatment (Fig. 2).

**Fig. 4.**
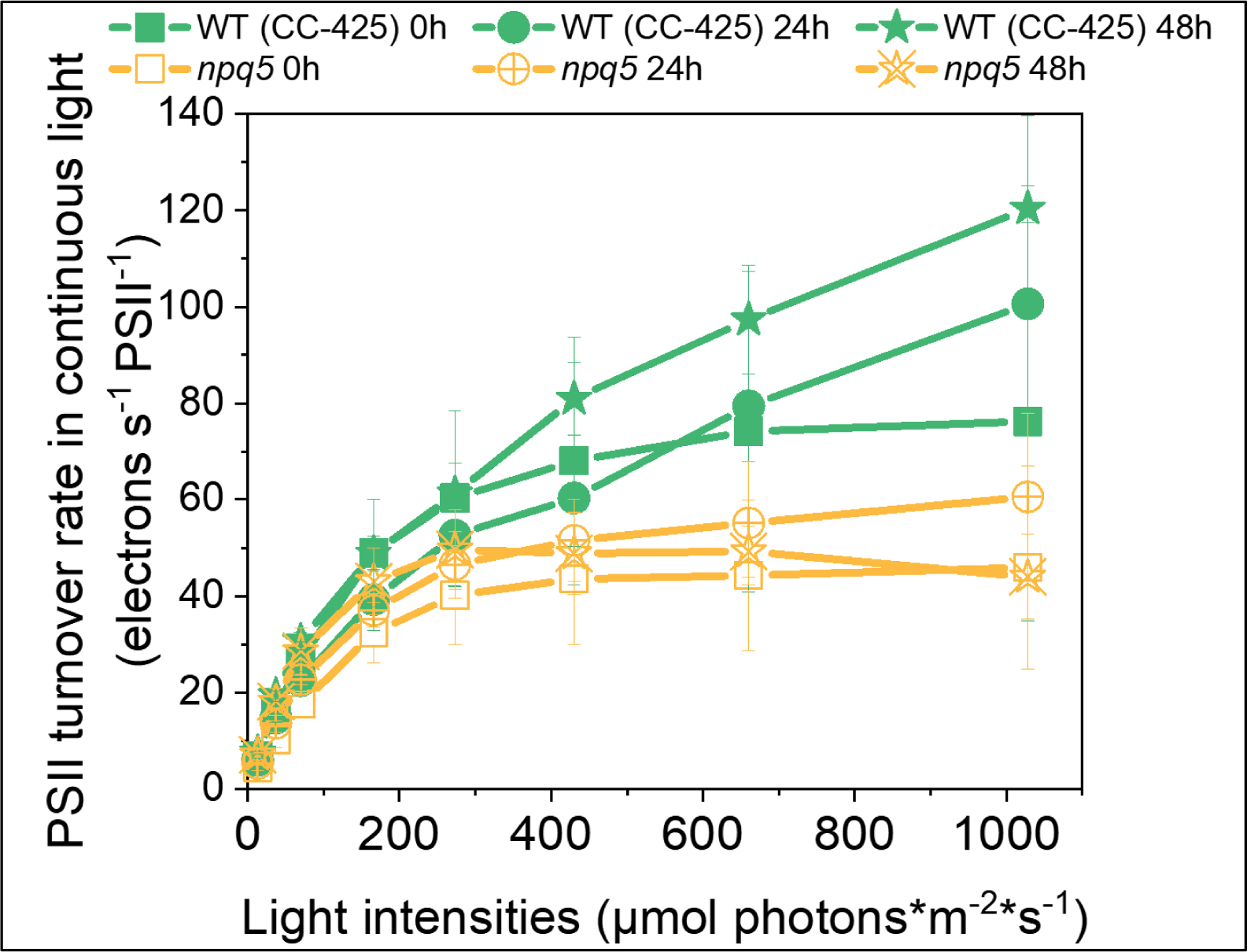
PSII turnover rate in continuous light. PSII turnover rate (y axis) was measured by multiplying Y(II) and maximal PSII rate at given light intensities (x axis). Data shown are mean ± SEM, n=3 biological replicas. See M&M for details about the calculation.

We also compared the abundance of the other major photosynthetic complexes in *npq5* and CC-425, using antibodies against ATP gamma subunit (ATPC), cytochrome f (Cyt f) and Rubisco Large subunit (Rubisco L). ATPC, Cyt f and Rubisco L showed only a slight difference between *npq5* and WT (CC-425) in both LL and HL (Fig. 3C-E, S2).

### Photoprotection capacity after high-light acclimation

Next, we examined the capacity of NPQ in the two strains. Since the amount of LHCSRs in the cells was shown to correlate with the level of NPQ (Bonente et al., 2012; Perozeni et al., 2018; Tian et al., 2019), we also quantified the LHCSR proteins during HL treatment.

NPQ was similarly low in both strains in LL and developed upon HL exposure only in WT (CC-425) (Fig. 5A). In *npq5*, the NPQ development was hampered, although there was a small increase after 48h in HL. Interestingly, LHCSR3 was expressed in both strains in HL: the LHCSR3/CP47 ratio was similar in WT (CC-425) and *npq5* at 24h HL, and increased further after 48h HL, but less in *npq5* than in the control (Fig. 5B). The LHCSR1/CP47 ratio was far lower in *npq5* than in WT (CC-425) at all time points (Fig. 5C). However, in HL-exposed cells LHCSR1 is present in a very low amount compared to LHCSR3 (Peers et al., 2009; Tian et al., 2019). To check how much the absence of LHCSR1 can affect NPQ, we examined the *lhcsr1* mutant (Allorent et al., 2016) and its control strain (WT 4A+) under the same light conditions. The absence of LHCSR1 had little effect on the NPQ level (Fig. S4) and it is thus unlikely to be the reason for the lower NPQ in *npq5*.

**Fig. 5.**
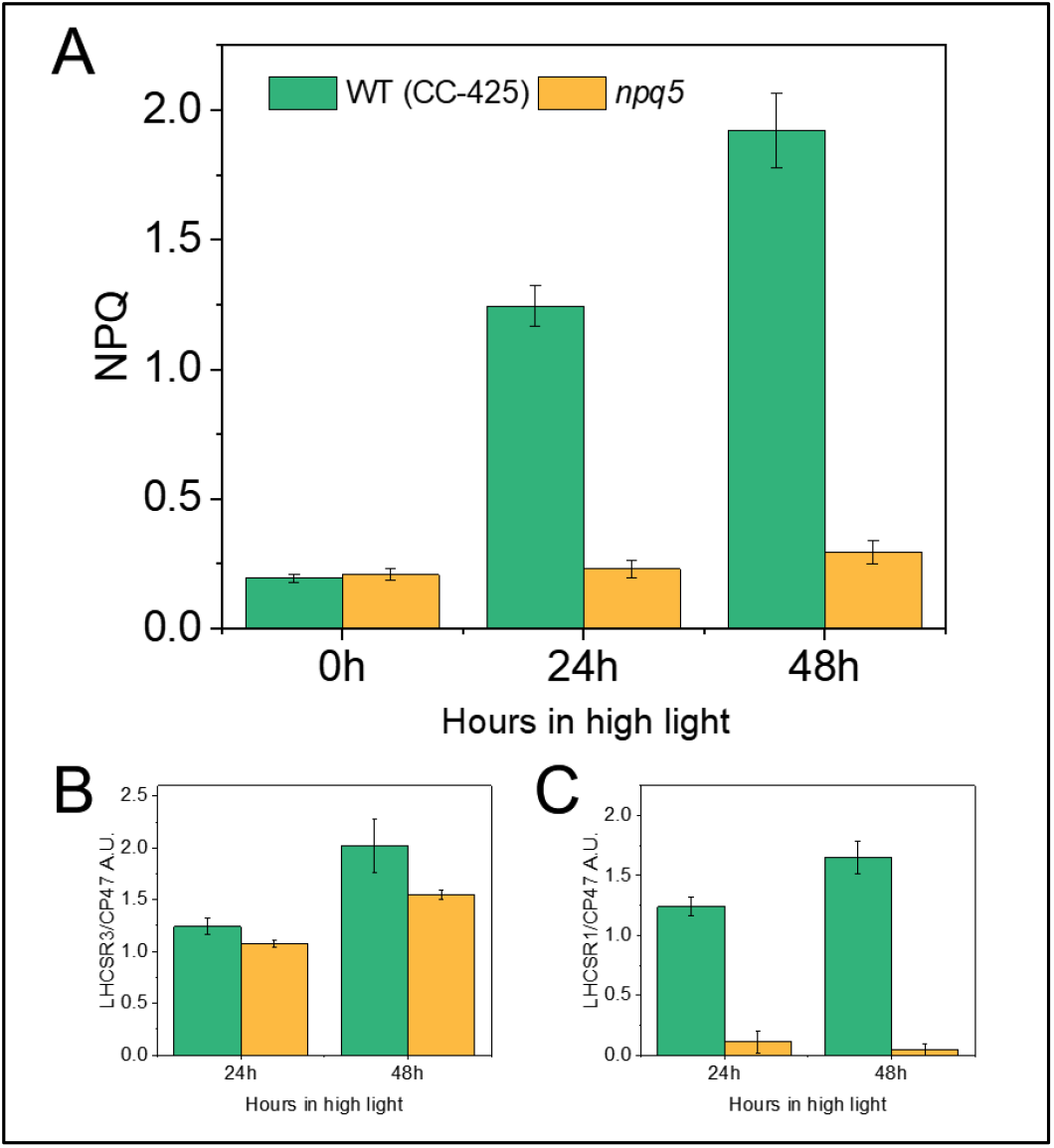
Maximum value for NPQ (A), LHCSR3/CP47 (B) and LHCSR1/CP47 (C) during HL treatment. Data shown are mean ± SEM, n=3 biological replicas. Densitometry data of each protein were normalized in WT (CC-425) at 24h (LHCSRs) or 0h (CP47) before calculating the ratio in (B-C).

### Is lumen acidification the bottleneck of NPQ development in *npq5 in vivo*?

NPQ is regulated not only by the amount of LHCSR3 but also by the acidification in the lumen. The lower PSII turnover rate in the mutant than in the control strain suggests a reduced capacity to create a proton gradient across the thylakoid membrane under light, which can be a reason for the low NPQ. To check this hypothesis, we bypassed the light-induced proton translocation to the lumen by adding acid to the cells. As shown previously, the addition of acetic acid to the cells induces stable, light-independent NPQ due to the LHCSR3 protonation (Tian et al., 2019). The quenching is reversible upon neutralization of the culture. We adopted this method to measure the maximal NPQ capacity at pH 5.5 in WT (CC-425) and *npq5*. In *npq5* the quenching was three times larger upon acid-induction than under light (Fig. 6A, S5A). However, the value at pH 5.5 was still far lower than that of the control strain at the same pH. To test if the cells reached the maximum NPQ capacity, we added a double volume of acetic acid. No further increase in NPQ was observed(Fig. S6). As shown by the scatterplot in figure 6B, the acid-induced NPQ maximum correlates with the amount of LHCSR3 per PSII (LHCSR3/CP47) in both strains, but is always lower in *npq5* than in CC-425. As an additional control, we also measured acid-induced NPQ in the *npq4* mutant, which does not contain LHCSR3, but contains WT level of LHCBM1 (Fig. S5B). At variance with *npq5*, the NPQ value remained very low in *npq4* also upon acidification of the lumen, in agreement with the role of LHCSR3 as pH sensor and NPQ effector.

**Fig. 6.**
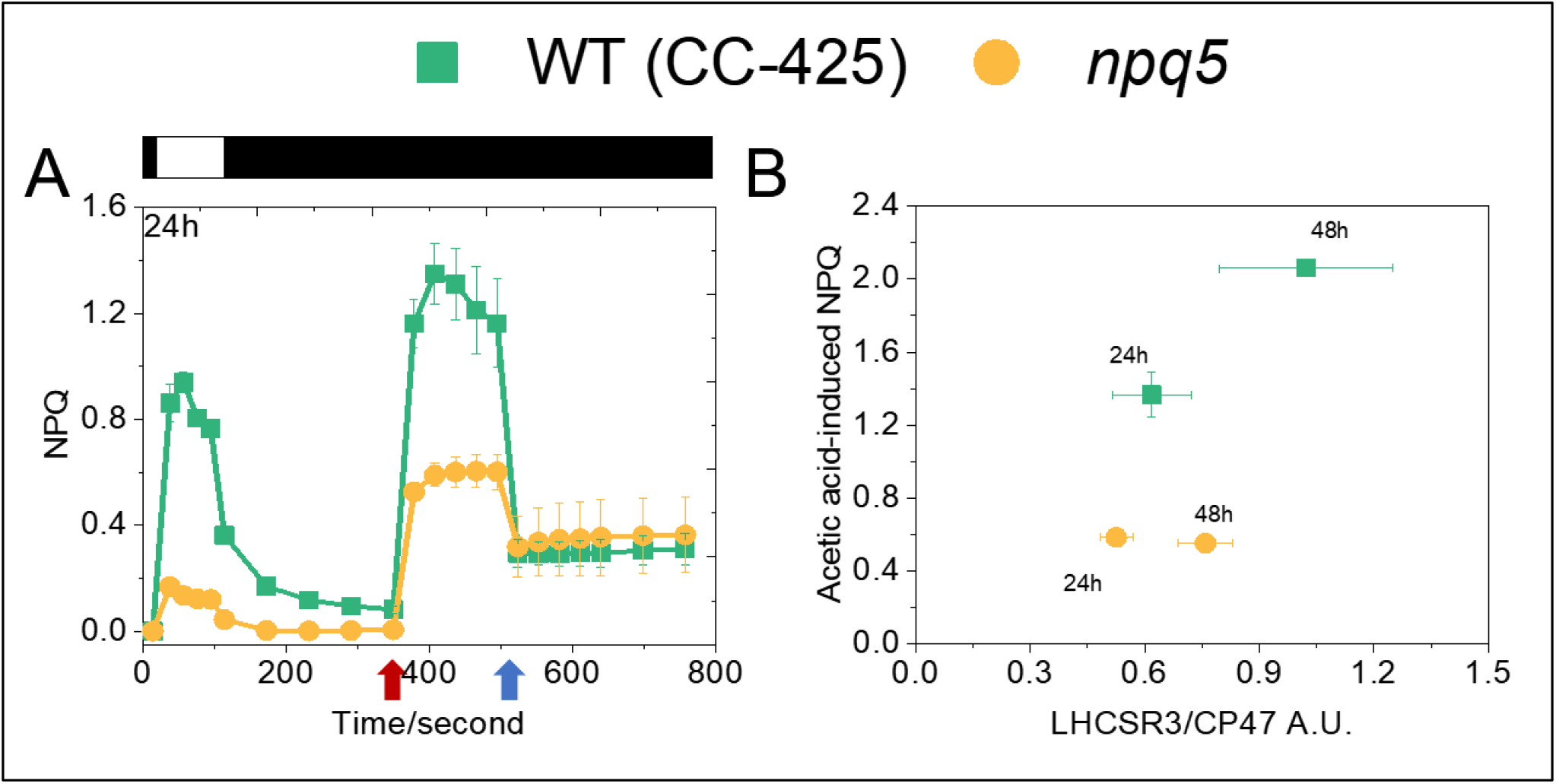
Comparison of light- and acid-induced NPQ in npq5 and WT (CC-425) (A) and correlation between the LHCSR3 content per core in the cells (expressed as LHCSR3/CP47 ratio) and acid-induced NPQ (B). The additions of 1 M acetic acid (decreases pH to 5.5) and 2 M KOH (neutralizes pH to 7.0) in (A) are indicated by red and blue arrows, respectively. Illumination (1500 μmol photons*m^-2^*s^-1^) and dark phases are indicated by white and black bars, respectively. Data shown are mean ± SEM, n=3 (CC-425); n=5 (npq5) biological replicas.

## Discussion

The main player in NPQ in *Chlamydomonas reinhartii* is LHCSR3 (Peers et al., 2009). This complex was shown to contain pigments (Bonente et al., 2011) and to switch from a light-harvesting to a quenched conformation in response to the protonation of its luminal residues (Liguori et al., 2013). LHCSR3 has thus been proposed to be both the pH sensor and the quencher in the membrane of *C. reinhardtii* (Tian et al. 2019). The absence of LHCSR3 in the cells (*npq4* mutant) completely abolishes the capacity of NPQ (Peers et al., 2009). A similar effect on NPQ was observed in the *npq5* mutant, which lacks LHCBM1 (Elrad et al., 2002), suggesting that this subunit is also involved in quenching. However, differently from LHCSR3 and LHCSR1, LHCBM1 is not a pH sensor and it is not able to switch from a light-harvesting to a quenched conformation in reponse to pH changes, as shown by both *in vitro* and *in vivo* experiments (Natali and Croce, 2015; Dinc et al., 2016) and confirmed here by the analysis of the *npq4* mutant. What is thus the reason for the substantial reduction in NPQ in the *npq5* mutant? The data shown in this work indicate that this effect is not due to a low amount of LHCSR3. It is also clear that despite in *npq5* the electron transfer chain works less effectively and the pH of the lumen is probably slightly higher than in the reference strain, this difference is not sufficient to explain the large difference in NPQ: acid-induced NPQ is still far smaller in the mutant than in the WT. Since upon 24h of HL (i) the amount of LHCSR3 is similar in *npq5* and its reference WT, (ii) PSII antenna size is similar in those strains, and (iii) when induced with acid, the lumen pH is the same, we can assume that the number of quenching centers (i.e. quenched LHCSR3) is also identical. What is thus the role of LHCBM1 in NPQ? One possibility is that the absence of LHCBM1 influences the connectivity between the quencher and the supercomplexes. LHCBM1 was shown to be part of trimer S, connecting this trimer with CP43 (Sheng et al., 2019). However, we can exclude that the absence of LHCBM1 alters the functional connectivity between the antenna and the core, because the maximal PSII rate in the mutant is very similar to that of the WT, correlating well with the LHCBM content. The maximal quantum yield of PSII (F_v_/F_m_) is also identical in mutant and WT, excluding differences in the presence of detached antenna complexes up to 24h of HL. It is thus likely that the absence of LHCBM1 influences the association of LHCSR3 with the supercomplex, thus limiting the quencher efficiency. Indeed, structural and spectroscopic data suggest that LHCSR3 interacts with the LHCBMs (Semchonok et al., 2017; Tian et al., 2019) and in particular, that LHCSR3 dimer is in direct contact with the LHCII-S trimer in the C_2_S_2_ PSII–LHCII supercomplex (Semchonok et al., 2017). A quenching rate of (150 ps)^-1^ was measured for Chlamydomonas WT cells in physiological conditions (Tian et al., 2019). This value is very similar to the overall trapping time in PSII supercomplexes (Caffarri et al., 2011), meaning that (i) LHCSR3 is well connected to PSII and (ii) is an efficient quencher (see (Croce and van Amerongen, 2020) for details on energy flow in the photosystems). In principle, LHCBM1 can influence both aspects. The interaction between LHCBM1 and LHCSR3 can create a more efficient quencher, for example, by stabilizing the quenched conformation of LHCSR3. The absence of LHCBM1 might also alter the association of LHCSR3 with the supercomplex, either because LHCBM1 is directly involved in the LHCSR3 docking or because its absence leads to the reorganization of the supercomplex, affecting LHCSR3 binding. As a result, less excitation energy reaches the quencher, explaining the NPQ phenotype. In this respect, it is interesting to mention that recent *in vitro* experiments have shown that LHCBM1 has a high tendency to interact with other LHCBMs. This property was suggested to depend on its charged N-terminus, which can help in protein-protein interactions (Kim et al., 2020), supposting the first hypothesis.

## Acknowledgments

This work is partially supported by the Human Frontiers Science Program (HRSP) grant RGP0005/2021. XL was supported by the Chinese Scholarship Council Fellowship 201606910042

## Supporting information

**Fig. S1.**
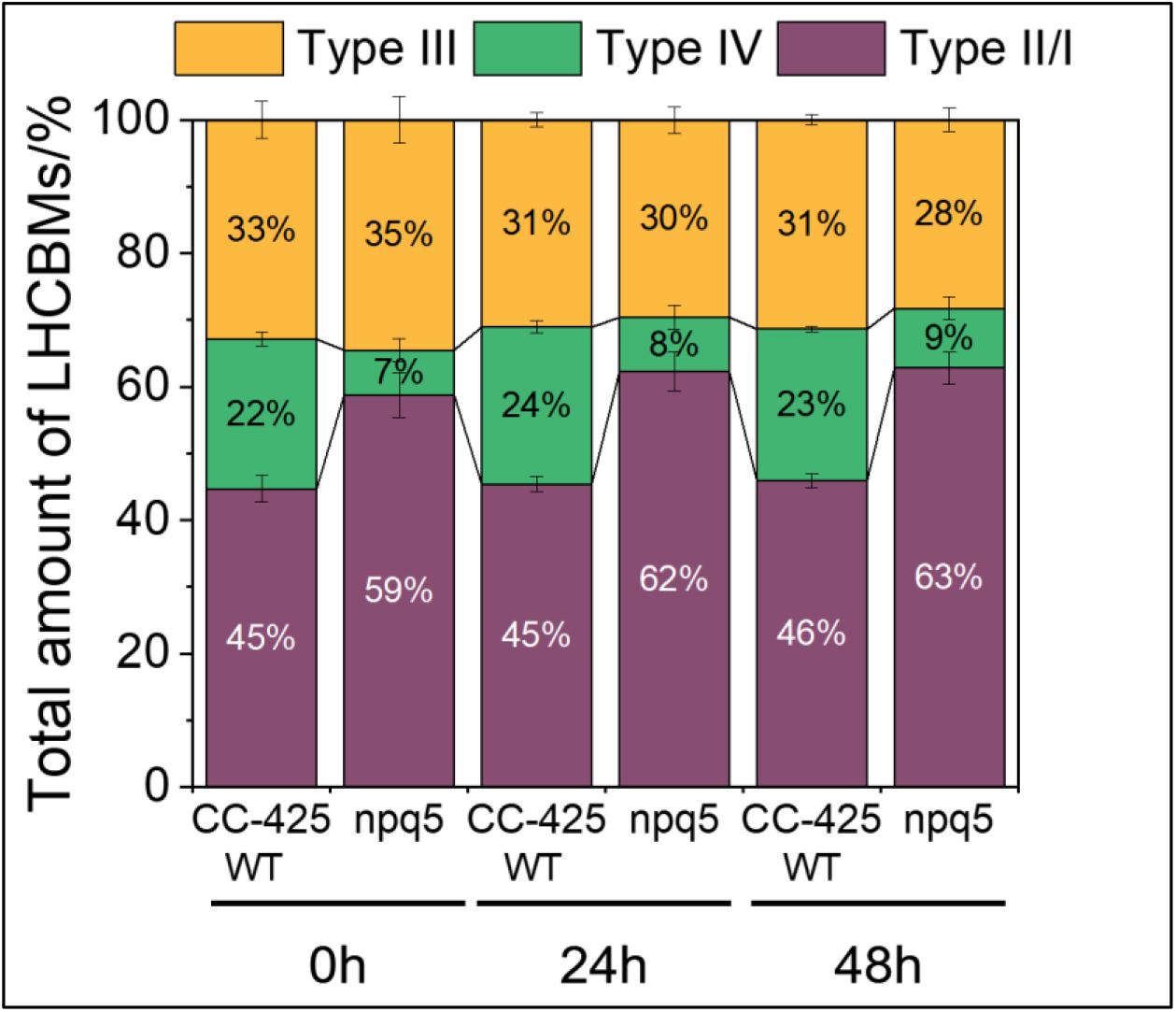
Quantification of LHCBM subtypes based on Immunoblotting during HL treatment in WT (CC-425) and npq5. Densitometry data of each subtype were normalized to the total amount of LHCBMs in each strain at each time point. Data shown are mean ± SEM, n=3 (WT CC-425) or 8 (npq5) biological replicas.

**Fig. S2.**
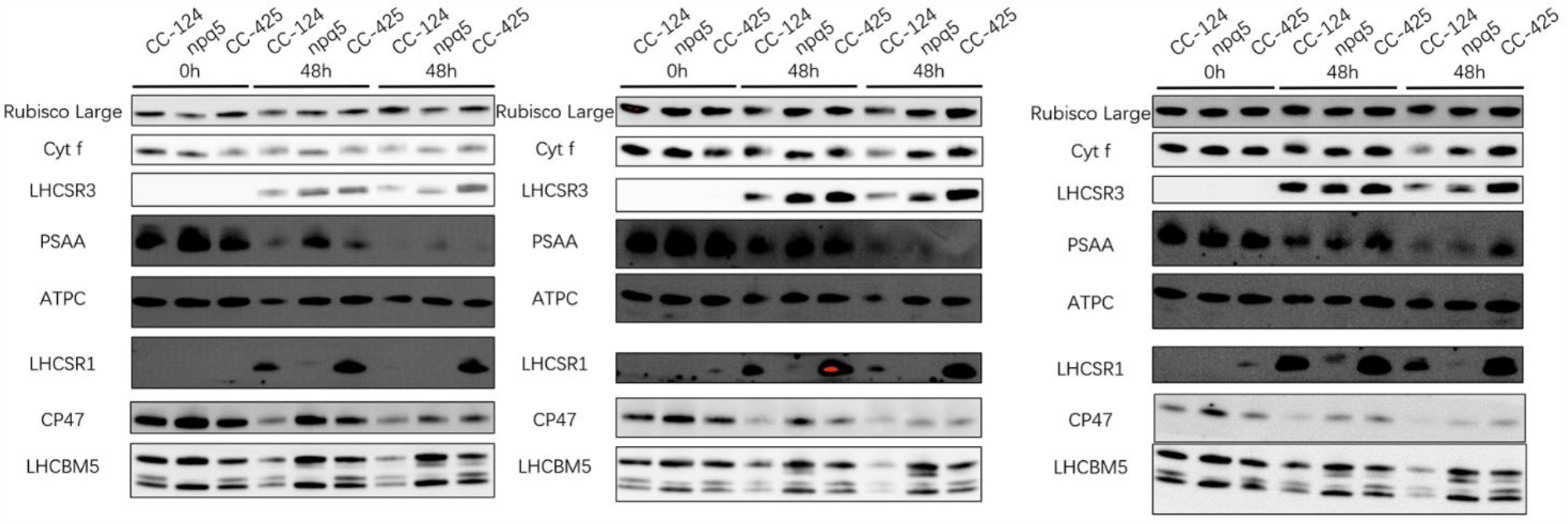
Immunoblots of photosynthetic apparatus during HL in WT (CC-124 and CC-425) and npq5. 5 μg total proteins were loaded in each well. Each panel shows one biological replica.

**Fig. S3.**
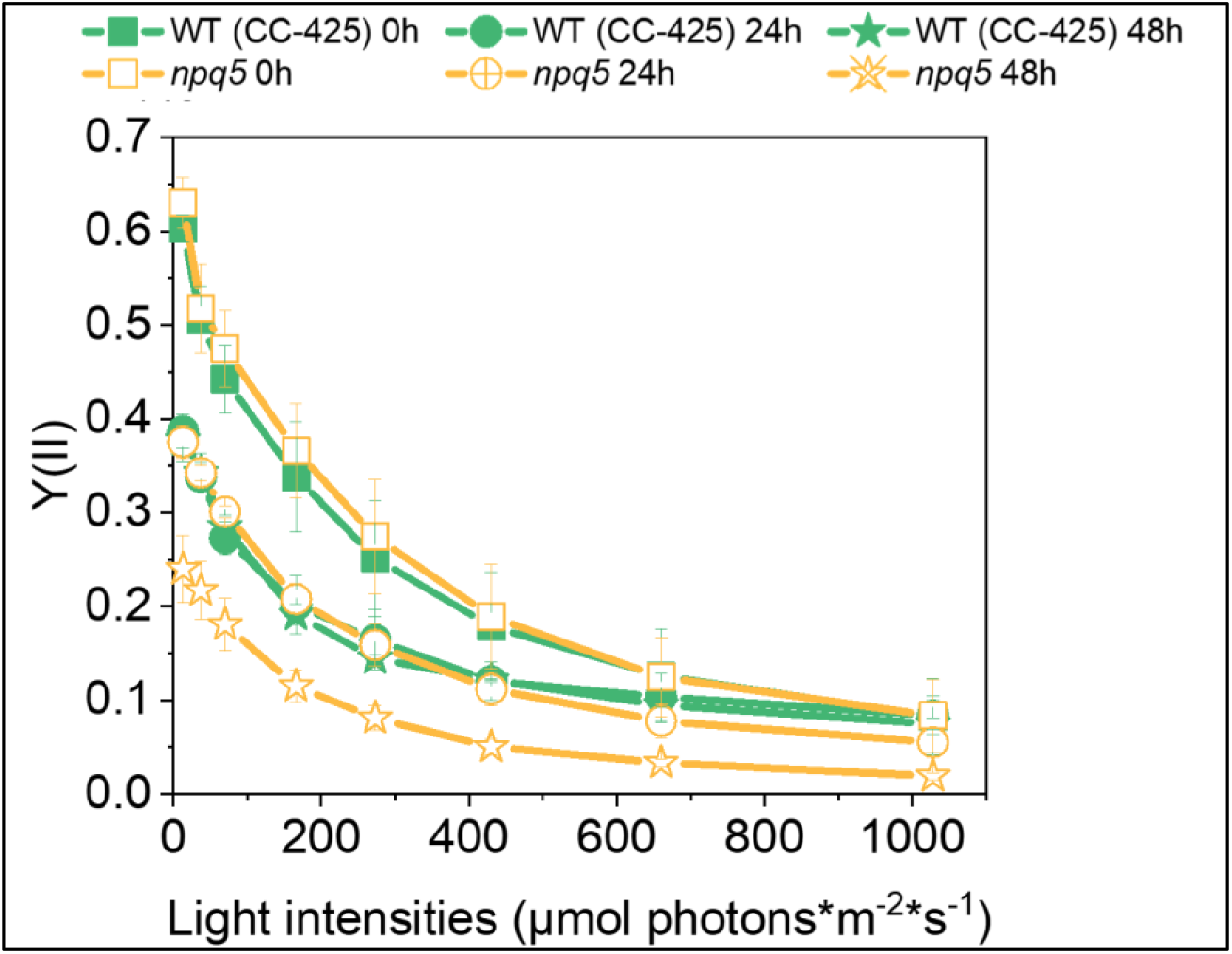
Y(II) was measured in DUAL-PAM 100 via light curve. Data shown are mean ± SEM, n=3 biological replicas.

**Fig. S4.**
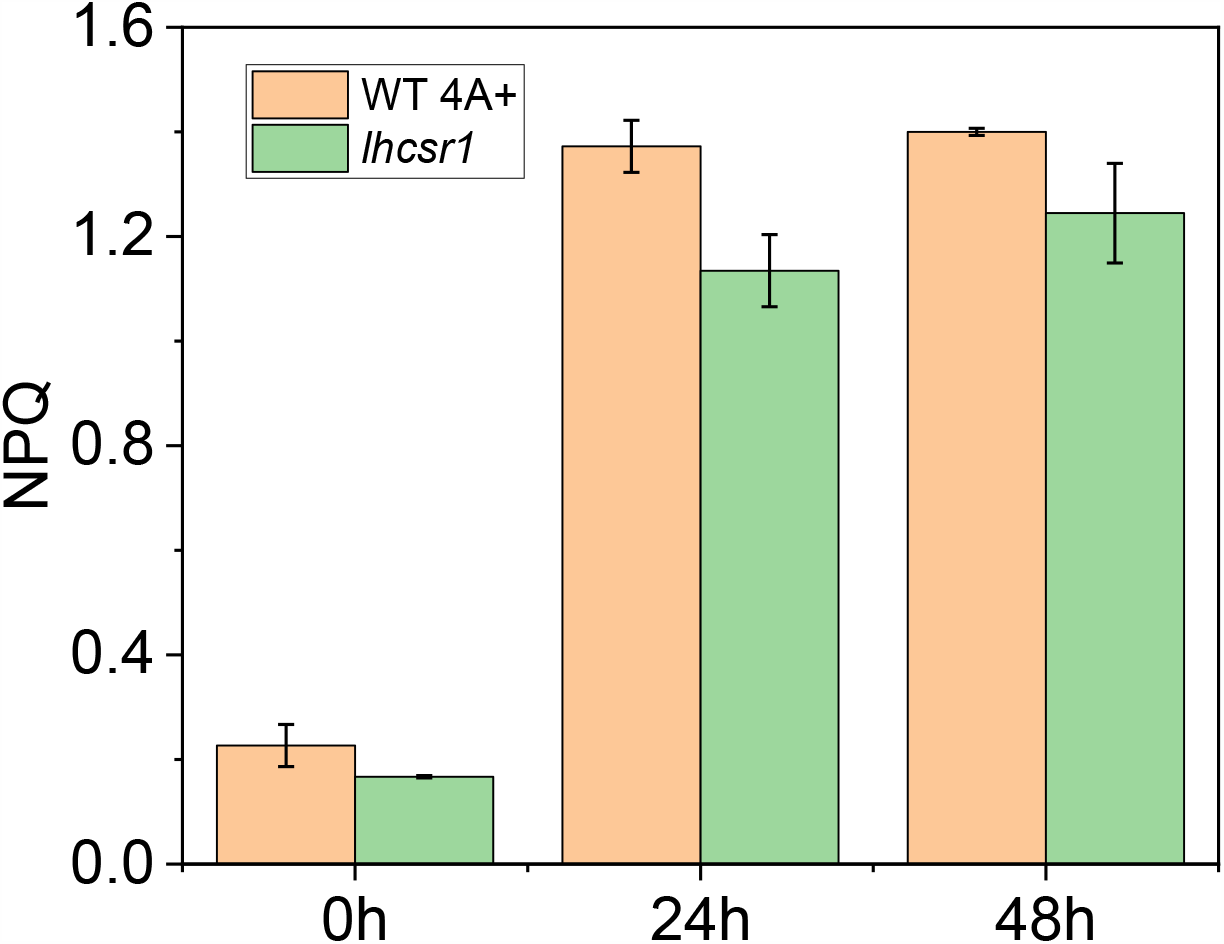
Maximum NPQ in WT (4A+) and lhcsr1 during HL treatment. The data are the result of two biological replicas.

**Fig. S5.**
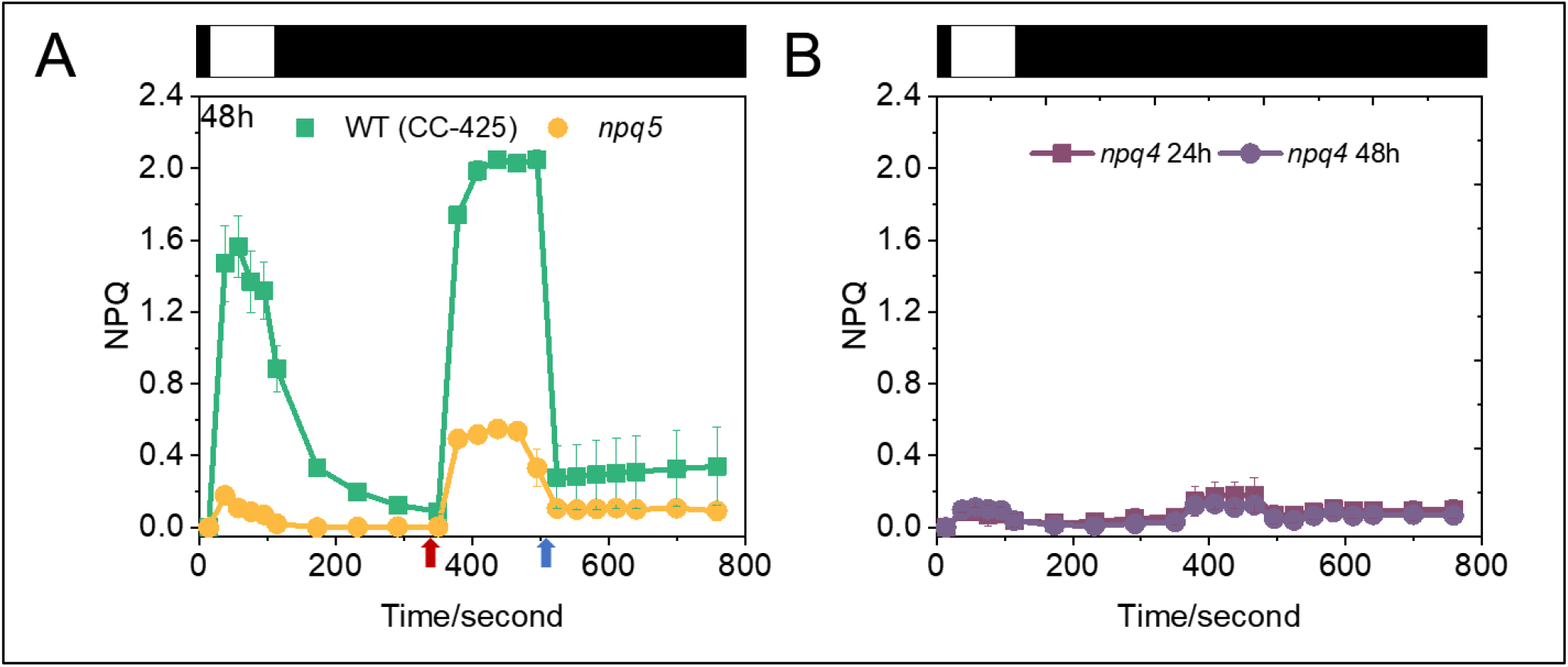
Comparison of light-induced and acid-induced NPQ in npq5 and WT (CC-425) at 48h (A) and npq4 during HL (B). The additions of 1 M acetic acid (decreases pH to 5.5) and 2 M KOH (neutralizes pH to 7.0) in (A) are indicated by red and blue arrows, respectively. Illumination (1500 μmol photons*m^-2^*s^-1^) and dark phases are indicated by white and black bars, respectively. Data shown are mean ± SEM, n=3 (WT CC-425) and n=5 (npq5) biological replicas in (A) and n=2 in (B).

**Fig. S6.**
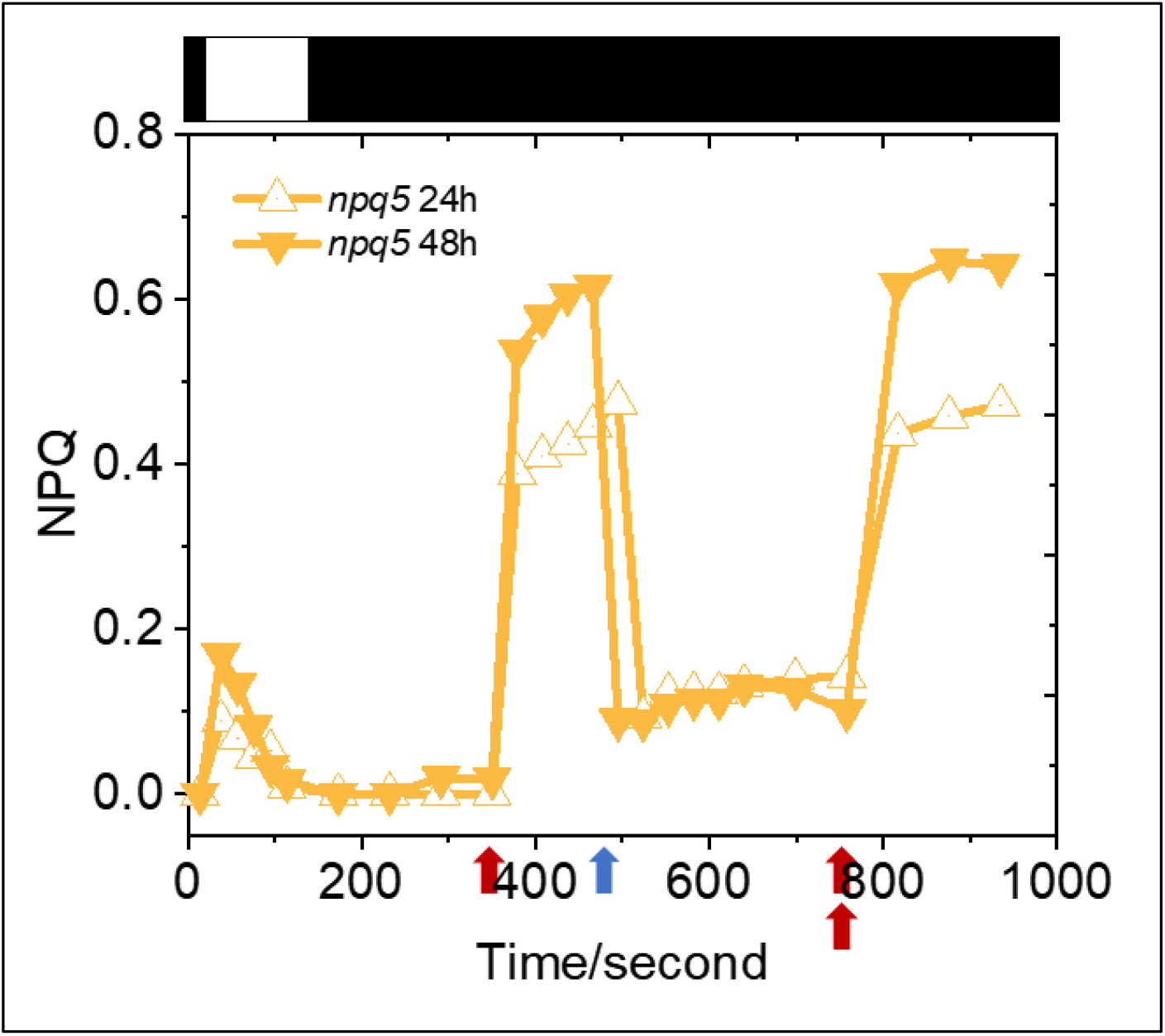
Double amount of 1 M acetic acid addition in npq5. The white and black bars indicate the illumination and recovery in darkness, respectively. The addition of acid or base is indicated by red or blue arrows, respectively. Two-red arrows indicate the double amount of acid added in the cell culture.

